# MeioBIOME: A snakemake workflow for the parallel analysis of meiofaunal genomes and host-associated bacteria/archaea

**DOI:** 10.64898/2026.07.23.740139

**Authors:** Alejandro De Santiago, Holly Bik

## Abstract

Microbes closely interact with every living organism, including meiofauna (i.e., microbial eukaryotes 38 μm – 1 mm in length), and influence the development, life cycle, and evolution of diverse metazoans. Together, meiofauna and their microbiomes, collectively referred to as the holobiont, underpin biogeochemical cycles and drive decomposition of organic matter. However, our understanding of the ecological and evolutionary dynamics of meiofauna microbiomes are limited, typically owed to low-resolution 16S rRNA surveys, which cannot accurately delineate bacterial taxa. Single-specimen holobiont sequencing can help overcome the limitations of metabarcoding approaches by 1) generating metagenome-assembled genomes (MAGs) of the host microbiome and 2) recovering host single-copy genes (SCGs) to phylogenetically confirm the identity of the host organism. However, most bioinformatics pipelines for the assembly of metagenomic datasets have been developed for the assembly of high-complexity microbial communities of bulk sediment or soil samples (and cannot be used for the assembly of host genomes), rely on co-assembly approaches (which collapses strain-level genomic information of bacterial taxa), and focus on binning either prokaryotic or eukaryotic taxa. Therefore, there is a tremendous need for a computational workflow for the dual analysis of host genomes and their microbiomes. Here, we developed MeioBIOME, a modular Snakemake pipeline for the reproducible analysis of holobiont metagenomes obtained from individually sequenced microbial metazoa. We analyze publicly available single-specimen metagenomics datasets to show the utility of MeioBIOME and recover host-associated symbiont MAGs and host SCGs. Additionally, we integrate state-of-the-art binning algorithms which generate more MAGs than the DOE Joint Genome Institute metagenomic pipeline. We anticipate that MeioBIOME will facilitate studies of phylosymbiosis by generating high-quality host genome skims (to build well-supported host phylogenetic trees) and host-associated prokaryotic MAGs obtained from single specimens.

## Introduction

Meiofauna (i.e., microbial eukaryotes 38 μm – 1 mm in length, such as nematodes, tardigrades, and copepods, etc.) are the most abundant and phylogenetically diverse group in ocean ecosystems that encompasses 23 animal phyla (Bonaglia et al., 2014; Danovaro et al., 2010; Laumer, 2024). Meiofauna harbor distinct microbiomes that differ from their environment (Boscaro et al., 2022), and there is substantial evidence that symbiosis is widespread across these phylogenetically diverse taxa (Sogin et al., 2021; Zeppilli et al., 2018). Seven animal phyla are known to harbor distinct lineages of chemosynthetic symbionts, including gutless flatworms (Platyhelminthes), oligochaetes (Annelida), and stilbonematids (Nematoda) (Bergin et al., 2018; Sogin et al., 2021; Woyke et al., 2006). These host-microbe interactions underlie key processes in the development, ecology, and evolution of these diverse animal taxa (Malard et al., 2021; Moreira et al., 2009), and it has led to the emergence of the “holobiont-hologenome” theory, which emphasizes that evolutionary pressures act on both the host and its associated microbial community (Madhusoodanan, 2019). Therefore, a major interest in microbial ecology is unraveling the community structure and metabolic potential of the bacterial assemblages associated with metazoan taxa to better understand evolutionary patterns of symbiosis of microbiomes and their host taxa.

Our knowledge of the biodiversity and evolutionary patterns of meiofauna microbiomes has largely been driven by high-throughput environmental DNA (eDNA) metabarcoding approaches using 16S rRNA single-marker surveys (Boscaro et al., 2022; Leasi et al., 2024; Schuelke et al., 2018). However, full and short-length 16S rRNA sequences, such as the V4 region commonly used for the high-throughput identification of bacterial taxa (Caporaso et al., 2012), provide limited taxonomic resolution as they are insufficient to delineate closely-related species (Strube, 2021). Additionally, full-length 16S rRNA gene trees are often in disagreement with whole-genome species trees and are inconsistent with average nucleotide identity (ANI - a method to assess genome relatedness and commonly used to delineate species boundaries of prokaryotic taxa; (Bartoš et al., 2024; Kiepas et al., 2024; Schloss, 2021)), indicating that single-marker genes cannot be used as a proxy to assess species diversity (and that 16S rRNA surveys likely underestimate the true microbial diversity of host and environmental microbiomes; (Durazzi et al., 2021)). Therefore, the use of single-marker genes may impact studies of phylosymbiosis due to the inability to distinguish between closely-related bacterial strains. For example, species boundaries of many bacterial taxa, such as *Pseudoalteromonas,* a bacterial genus with a cosmopolitan distribution in marine environments that is commonly found as a core member of marine invertebrate microbiomes (Bellec et al., 2019; Ohdera et al., 2023; Pereira et al., 2023; Shan et al., 2019), cannot be delineated using the full-length 16S rRNA gene due to low sequence divergence among species, intragenomic variation within each genome, and multiple gene variants that are shared across closely-related taxa (Strube, 2021). Despite *Pseudoalteromonas* being recognized as a member of the core microbiome of several marine invertebrates (and underpinning the development and life cycle of diverse metazoan taxa; (Ma et al., 2014; Malter et al., 2025; Ohdera et al., 2023)), many 16S rRNA studies have found conflicting evidence of phylosymbiosis among marine invertebrates, with the largest study of meiofaunal microbiomes to date finding no evidence of phylosymbiosis across 21 invertebrate phyla (Boscaro et al., 2022), while other taxon-focused 16S rRNA studies have concluded that phylosymbiosis does exist, such as nemertean worms (Leasi et al., 2024) and brachyuran crabs (Tsang et al., 2024). However, using whole-genome data, De Santiago et al. (2026) constructed a pangenome of free-living and invertebrate-associated *Pseudoalteromonas* and found cophylogenetic signals between *Pseudoalteromonas* and three invertebrate phyla (Mollusca, Nematoda, Cnidaria). Conflicting results between 16S rRNA surveys and whole-genome datasets indicate that the recovery of whole-genome data of host-associated bacteria and single-copy orthologous genes of the host organism (to build well-supported phylogenetic species trees; (Alam et al., 2025)) can help recover signals of phylosymbiosis.

Despite the importance of whole-genome data, meiofauna have largely been ignored in whole-genome sequencing initiatives due to a combination of low DNA biomass of individuals (microbial metazoa can contain <1,000 somatic cells; (Cohen-Fix & Askjaer, 2017)), and high levels of cryptic diversity (which prevents pooling a large number of individuals for sequencing since it can affect assembly quality due to increased genomic heterogeneity; (Bik, 2019)). This roadblock has affected our ability to gain scientific insights into the ecology and evolution of meiofauna and their microbiomes. A recent genome census estimated that ∼50% of invertebrate-associated prokaryotic genera lack any genomic representation (including isolate genomes or MAGs; (Wu et al., 2025)), despite the high estimated prevalence of symbiosis among invertebrates (with studies of insect symbiosis estimating that 20% of insect species harbor intracellular symbionts; (Douglas, 2011; Venn et al., 2008)). In stark contrast, almost no human-associated bacterial genera lack a genomic representative, which has been largely driven by the dedicated genomic sequencing efforts of the Human Microbiome Project (Turnbaugh et al., 2007). Recent advances in molecular biology have improved our ability to extract and amplify High Molecular Weight (HMW) DNA from meiofauna (Qing et al., 2025; Roberts et al., 2024). For example, Roberts et al. (2024) found that whole-genome amplification (WGA) methods can be used to generate enough DNA template for PacBio HiFi long-read sequencing and resulted in the first complete genome assembly of a ∼250 µm Gastrotrich (*Lepidodermella squamata*). Furthermore, these enzymatic methods have been used to generate host genome skim data (single-copy genes that can be used to construct well-supported phylogenomic trees) to expand our current knowledge of the evolution of marine nematodes (Qing et al., 2025) and can also amplify bacterial DNA as it has been used to quantify the phylogenomic diversity, abundance, and prevalence of *Pseudoalteromonas* in nematode microbiomes (De Santiago et al., 2026).

There have been substantial efforts to develop comprehensive bioinformatics algorithms and pipelines to assemble MAGs from metagenomic datasets (Churcheward et al., 2022; Hofmeyr et al., 2020; Kang et al., 2019; Kieser et al., 2020; Wang et al., 2024; Zou et al., 2025), which have led to the discovery of novel difficult-to-culture prokaryotic lineages and have transformed our knowledge of the tree of life (Hug et al., 2016; Nayfach et al., 2021; Villada et al., 2025; Wu et al., 2025). However, many of these tools have primarily focused on either binning only prokaryotic (Churcheward et al., 2022; Kieser et al., 2020) or only eukaryotic MAGs (with a primary focus on binning protist genomes, such as the EukHeist pipeline; (Alexander et al., 2023)) from bulk sediment or soils. Additionally, they often rely on the co-assembly of related samples (Hofmeyr et al., 2020) to improve the recovery of closely-related MAGs, which can affect cophylogenetic studies by collapsing strain-level information (Vosloo et al., 2021).

Pipelines designed for the recovery of host-associated MAGs often neglect host genome data (Bai et al., 2025; Foo et al., 2023), despite host phylogenies being necessary to understand the ecology and evolution of symbiont taxa. Additionally, state-of-the-art binning software, such as COMEBin (Wang et al., 2024) and CompleteBin (Zou et al., 2025), which implement novel deep-learning binning algorithms, have been shown to increase the recovery of MAGs in real host-associated samples and in simulated datasets with high strain-level diversity (Wang et al., 2024; Zou et al., 2025), but they have yet to be integrated in modern bioinformatics pipelines. Therefore, there is a need for updated computational workflows designed for the parallel analysis of meiofaunal metagenomic datasets, where shotgun metagenomic sequencing is required to 1) identify the taxonomic classification and phylogenetic placement of host taxa and 2) assemble novel host-associated prokaryotic MAGs.

Here, we present MeioBIOME (Meiofauna microBiome), a Snakemake bioinformatics workflow (Köster & Rahmann, 2012) for the reproducible and parallelized recovery of host genome skim data and host-associated bacterial and archaeal MAGs. MeioBIOME integrates reliable open-source computational tools, including two novel machine-learning binning algorithms that utilize a pretrained deep-learning language model with dynamic contrastive learning (i.e., CompleteBin and COMEBin) to increase the recovery of near-complete bacterial MAGs without co-assembling of samples. We show the utility of MeioBIOME by analyzing a publicly available single-nematode metagenomic dataset generated using whole-genome amplification methods (De Santiago et al., 2026) and show that we can recover a substantial number of lineage-specific single-copy ortholog genes of the host taxa. Additionally, the pipeline recovers more MAGs than the commonly used DOE JGI Integrated Microbial Genomes (IMG) binning pipeline. Therefore, MeioBIOME will help facilitate studies of phylosymbiosis by assembling genomic data of taxa that are often neglected from sequencing initiatives (Bik, 2019), which can help answer fundamental questions in meiofauna research (Martínez et al., 2025) and promote the discovery of new lineages of life (Wu et al., 2025).

## Design and Implementation

### Implementation and Input Files

The primary input for the MeioBIOME pipeline v1.0 is a directory that contains raw non-interleaved paired-end sequence files using the following specific, yet simple, naming convention: *samplename_R[1/2].fastq.gz* (**Figure 1**). In addition, a configuration file allows for the customization of certain parameters throughout the workflow, while the cluster configuration file allows the pipeline to be easily parallelized and run on a high-performance computing cluster (HPCC; (Mölder et al., 2021)). At each step, the pipeline utilizes Conda to install the necessary open-source computational tools (**Table S1**). The pipeline can be easily run using five commands in the following order:

1. *snakemake --until **reads_quality_trim** --cluster-config cluster.yaml --snakefile snakefile*
2. *snakemake --until **metagenome_assembly** --cluster-config cluster.yaml --snakefile snakefile*
3. *snakemake --until **host_genome_skim** --cluster-config cluster.yaml --snakefile snakefile*
4. *snakemake --until **microbiome_bins** --cluster-config cluster.yaml --snakefile snakefile*
5. *snakemake --until **microbiome_bins_quality** --cluster-config cluster.yaml --snakefile snakefile*
6. *snakemake --until **microbiome_bins_quant** --cluster-config cluster.yaml --snakefile snakefile*

**Figure 1.**
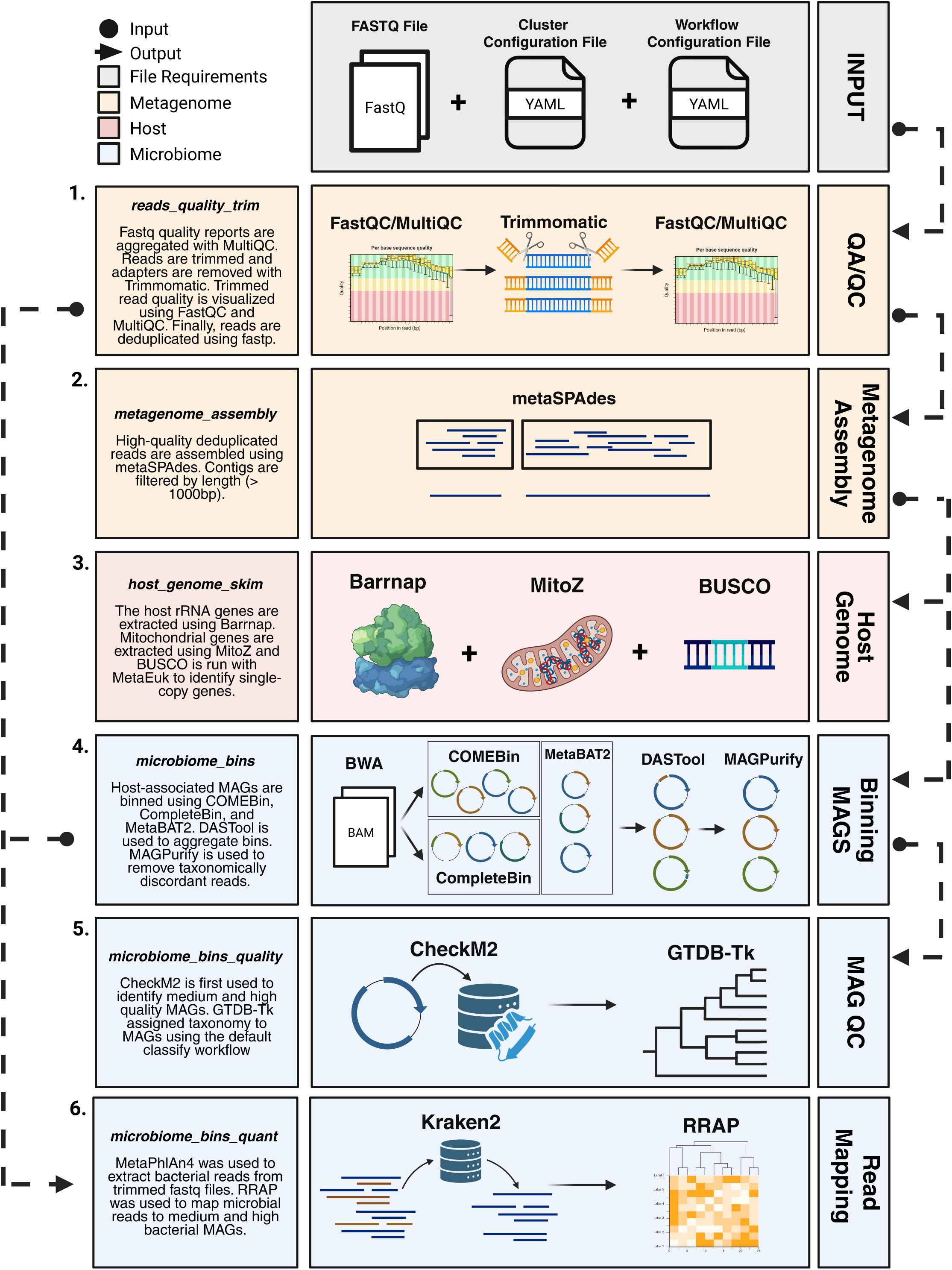
Schematic of the workflow implemented in MeioBIOME. There are two files required to run the metagenomic workflow in addition to the fastq files: a cluster and a workflow configuration file. Fastq files undergo 1) adaptor removal, quality trimming, and 2) metagenomic assembly. Only contigs larger than 1,000 bp are kept for further analysis. Afterwards, the metagenome assemblies undergo 3) host-skimming (identification of single-ortholog genes and mitochondrial genome) and 4) binning of metagenome-assembled genomes (MAGs). Finally, 5) MAG quality is assessed using CheckM2 and taxonomically identified using GTBD-Tk workflow. Created in https://BioRender.com

Each workflow step is tracked with log files describing the command, input, output, and resource requirements, and records errors that arise. The tools included in the MeioBIOME pipeline (which should also be cited when using this tool) are extensively documented in **Table S1**. The final outputs include genome skims of the host organism, host-associated prokaryote metagenome-assembled genomes (MAGs), and summary files (**Figure S1**).

### Sequence Quality Control and Metagenomic Assembly (Steps 1 and 2)

Sequence quality and adapter content are first analyzed using FastQC (Andrews & Others, 2010), and individual reports are aggregated and visualized using MultiQC (Ewels et al., 2016). Subsequently, reads undergo adapter removal and quality trimming using the Trimmomatic software (Bolger et al., 2014). Reads less than 55 bps, the maximum k-mer length implemented using default metaSPAdes (Nurk et al., 2017) parameters, are removed. The quality-controlled sequences are once again analyzed using FastQC and MultiQC, allowing users to easily identify whether the reads were processed appropriately or if they require different trimming parameters. Adapter sequences and Trimmomatic parameters can be easily updated by modifying the config file. Next, duplicate sequences are removed using Fastp (Chen et al., 2018) to reduce the complexity of the dataset and remove technical errors, which is known to reduce computational resources and runtime and increase contig continuity.

High-quality sequences are then assembled into scaffolds using metaSPAdes (Nurk et al., 2017). To reduce the computational workload during host genome skimming and metagenomic binning, scaffolds less than 1,000bp are removed using a Python script provided by COMEBin (Wang et al., 2024).

### Host Genome Skimming (Step 3)

First, BUSCO (Manni et al., 2021) is first run in batch mode using genome mode, using the OrthoDB v12 (Tegenfeldt et al., 2025) lineage-specific protein reference database and the MetaEuk (Levy Karin et al., 2020) protein predictor to identify and extract phylum-specific single-copy genes. Second, MitoZ (Meng et al., 2019) is run to extract and annotate mitochondrial contigs that were assembled by metaSPAdes. Finally, Barrnap (https://github.com/tseemann/barrnap) is used to extract the 18S, 5.8S, and 28S ribosomal RNA (rRNA) genes of the host organism.

### Assembling Microbial MAGs (Steps 4)

BWA (Li, 2013) and SamTools (Danecek et al., 2021) are used to map the trimmed the deduplicated short-reads to the metagenome-assembled scaffolds (>1,000bp) to quantify the abundance of each contig in the dataset. This read abundance information is used by three binning algorithms: CompleteBin (Zou et al., 2025), COMEBin (Wang et al., 2024), and MetaBAT2 (Kang et al., 2019) to bin assembled reads into metagenome-assembled genomes (MAGs). COMEBin implements a contrastive multiview-representation learning to help improve the assembly quality and has been shown to improve the recovery of novel lineages with small genomes that lack most universal markers (i.e., bacterial lineages within the Candidate Phyla Radiation). CompleteBin applies a dynamic learning algorithm with a novel pre-trained deep language model that implements the Leiden algorithm to recluster bins with high contamination (>10%) or remove short contigs with redundant single-copy genes containing single in bins with moderate contamination (<5%). DASTool (Sieber et al., 2018) is run on the binning outputs using a score_threshold of 0 to dereplicate bins without filtering for completion (number of expected single-copy genes recovered using a bacteria or archaea specific dataset) or contamination (number of duplicated single-copy genes). Afterwards, a Python script is used to keep bins that have a minimum of 50% completeness and a maximum of 10% contamination (according to a set of bacterial and archaeal single-copy genes employed by DASTool). The completion and contamination filters can be easily modified by updating the workflow config file. Each MAG undergoes QA/QC using four distinct modules implemented in MAGpurify (Nayfach et al., 2019): taxonomically discordant contigs are removed using the *phylo-markers* and *clade-markers* modules, and contigs with outlier GC content and known human or phi-x contaminant are removed with the *gc*-*content and known-contam* modules.

### Assess Quality and Quantifying Abundance of MAGs (Step 5 and 6)

CheckM2 (Chklovski et al., 2023), which uses several genomic features, such as the number of coding sequences and amino acid counts, in addition to single-copy genes, is used to assess MAG quality. Bins that are classified as either high-quality (>90% completion and <5% contamination), medium-quality (>50% completion and <10% contamination), or low-quality (<50% completion or >10% contamination) per community standards (Bowers et al., 2017) were kept for further analysis. The GTDB-Tk v2.4.1 classify workflow (Chaumeil et al., 2022), using the flag *--skip-ani-screen* to skip the pre-filter fastANI screen (Jain et al., 2018), is used to taxonomically identify each bacterial MAG. The databases used for QA/QC and taxonomic ID can be easily updated by specifying the file path to the reference database in the config file.

Finally, Kraken2 (Wood et al., 2019) is used to extract bacterial reads from the quality-filtered and deduplicated short reads from MeioBIOME step 1 (*read_quality_trim* command). The bacterial reads were mapped to the high- and medium-quality bacterial bins using RRAP (Kojima et al., 2022), and their abundance is quantified in Reads Per Kilobase of transcript per Million mapped reads (RPKM).

## Results

### Validating pipeline using publicly available metagenomic datasets of individual nematodes

We used 21 publicly available datasets (De Santiago et al., 2026) generated using single-worm shotgun metagenomic sequencing of four nematode families (Thoracostomopsidae, Oncholaimidae, Sphaerolaimidae, and Leptosomatidae) to show the utility of the pipeline (**Table S2**). These samples were processed using three different methods: 1) Frozen EZNA, 2) Live EZNA, and 3) Live RepliG. Frozen EZNA samples were previously frozen in bulk and underwent DNA extraction using the Omega EZNA DNA Isolation Kit. Live samples were stored in TL Buffer before undergoing DNA extraction using the Omega EZNA DNA Isolation Kit. Finally, the Live RepliG were live specimens that were stored in REPLI-g Single Cell Cryo-protect Reagent (QIAGEN) after isolating them from marine sediments. Worms in REPLI-g Single Cell Cryo-protect Reagent (QIAGEN) underwent DNA lysis and multiple displacement amplification (MDA) using the REPLI-g Advanced DNA Single Cell Kit (QIAGEN). Samples were processed using the default settings for MeioBIOME and were simultaneously processed using the JGI IMG workflow. The recovered MAGs were further placed in phylogenetic trees using GToTree (Lee, 2019), and the data (recovery of SCGs and MAG quality) was summarized in RStudio.

BUSCOs were recovered using the nematode-lineage specific dataset from the OrthoDB v12 and include 597 single-copy genes identified from 79 genomes (which are largely biased to agricultural and terrestrial parasitic nematode species and thus the “true” set of SCGs likely varies; **Figure 2A and Table S2**). Aside from a single sample (Oncholaimidae), MeioBIOME recovered <18 (3.02%) nematode-specific BUSCOs from samples that were previously frozen prior to isolation and DNA extraction (Leptosomatidae and Oncholaimidae). In contrast, 97.15%-99.50% of nematode-specific BUSCOs were recovered from live EZNA and RepliG Oncholaimidae samples (Complete: 18-135; Duplicate: 37-201; Fragmented: 245-506), and roughly 25.80%-41.37% of BUSCOs were recovered from the RepliG Thoracostomopsidae samples (Complete: 27-101; Duplicate: 2-25; Fragmented: 104-149). Finally, 53.10%-58.29% of BUSCOs were recovered from the live E.Z.N.A. Sphaerolaimidae samples (Complete: 95-118; Duplicate: 3-11; Fragmented: 206-225). This finding is consistent with a previous phylogenetic study of marine nematodes that found that recovery of metazoan-specific BUSCOs ranged from 50%-99% from samples that were immediately stored in RepliG storage buffer (Qing et al., 2025). Considering that the nematode BUSCO dataset is composed of mostly parasitic and agriculturally important taxa (and is therefore bias against free-living and marine species), the “true” set of nematode SCGs may vary. However, more experiments are needed to confirm these results.

**Figure 2.**
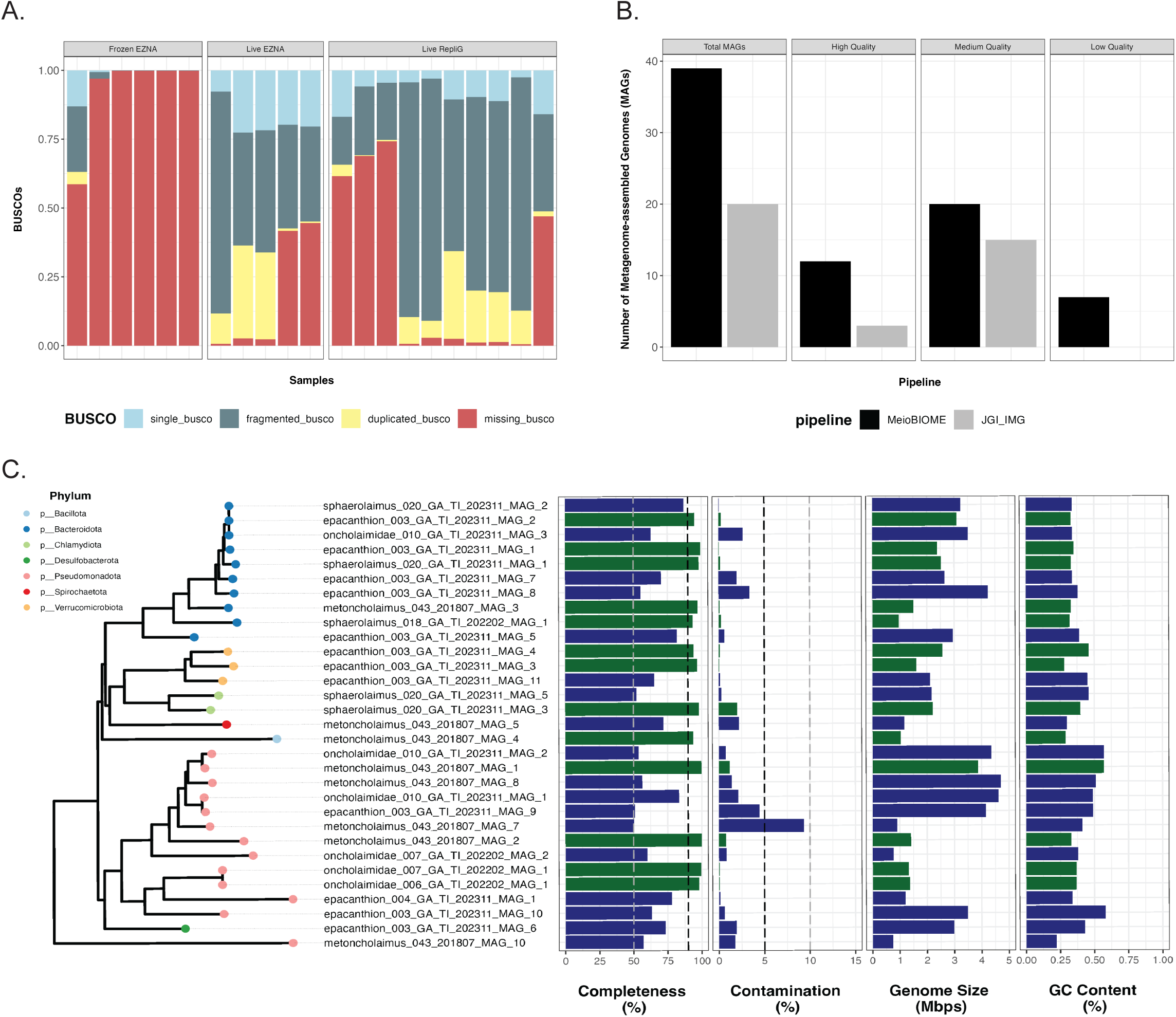
Recovery of Genome Skims and Host-associated MAGs of 21 publicly available single-worm metagenomic datasets. A) The recovery of single-copy ortholog genes in each metagenomic dataset. B) Recovery of metagenome-assembled genomes (MAGs) compared to the JGI IMG pipeline (last updated July 25th, 2025). C) Summary of the completion, contamination, genome size, and GC content of the recovered MAGs. The midpoint-rooted maximum likelihood phylogenetic tree was built using 74 bacteria-specific single-copy genes using GToTree (Lee 2019). The color of the bars indicates whether the MAG is high (green) or medium (blue) quality.

The host-associated bacterial MAGs were recovered using MeioBIOME default parameters. A total of 40 MAGs were recovered from 8 samples (2 were classified as common contaminants and were removed from further analysis; **Figure 2B and Table S3**). Of the 38 host-associated MAGs, 13 of the MAGs were classified as high-quality (>90% completion and <5% contamination), and 16 MAGs were medium-quality (>50% completion and <10% contamination). 9 MAGs were low-quality (<50% completion or >10% contamination) and were not included in the phylogenetic analysis due to either high contamination or low recovery of single-copy genes. Additionally, compared to the new JGI IMG Binning pipeline (last updated July 2025; (Clum et al., 2021)), we recover nearly 2X more total host-associated MAGs, including 4X and 1.3X more high and medium-quality MAGs, respectively (**Figure 2B**). Only one single MAG (*Pseudoalteromonas undina*) had more than 5% contamination (**Figure 2C**), which can likely be attributed to the high strain-level diversity of nematode samples (De Santiago et al., 2026). De Santiago et al. (2026) previously showed that the Oncholaimidae sample harbors a high abundance of a diverse number of *Pseudoalteromonas* species, which impact the recovery of high and medium-quality MAGs. It should be noted that *P. undina* was previously isolated from a marine nematode belonging to the same taxonomic family as this sample (De Santiago et al., 2025). Among the 38 host-associated MAGs we recovered, most of the MAGs were classified as *Bacteroidota* (10), a diverse group of bacteria that degrade complex carbohydrates, and *Pseudomonadota* (15). Among the 38 host-associated MAGs we recovered, there were 4 *Rickettsiales* and 3 *Chlamydiota* genomes (**Table S3**). The four *Rickettsiales* MAGs were recovered from three Oncholaimidae and one Thoracostomopsidae samples, and two were identified as *Ca. Lariskella* (G964019805), which was recently confirmed to infect the reproductive organs of female nematodes (Hagenbeek et al. 2025). The *Chlamydiota* MAGs were recovered from a single sample (Xyalidae). Two of the *Chlamydiota* MAGs were classified as *Simkaniaceae* (an obligate intracellular bacterium previously associated with gutless oligochaetes (Kleiner et al. 2012; Köstlbacher et al. 2021)) and SLFE01 (closely-related to MAGs isolated from a hypersaline lake and a marine sediment core; **Table S3**).

## Availability and Future Directions

The source code of MeioBIOME is available on GitHub (https://github.com/BikLab/meioBIOME-metagenomics). MeioBIOME provides a standardized and reproducible workflow that has been shown to recover host genome skim data (18S rRNA gene, partial mitochondrial genomes, and single-copy orthologs) and host-associated bacterial MAGs, including known endosymbionts of marine invertebrates. Finally, due to the modular framework of Snakemake, future updates will focus on extending the usability of MeioBIOME by integrating long-read PacBio and Nanopore sequencing and co-binning of samples to help recover more high and medium-quality MAGs without co-assembly of samples.

## Acknowledgement

Funding for this study was provided by the Gordon and Betty Moore Foundation (Symbiosis in Aquatic Systems Initiative, grant #9326), a National Science Foundation CAREER award (DEB-2144304), and an Antarctic Program award (OPP-2132641) to HMB at UGA. Research support for ADS was provided by the University of Georgia Research Foundation and the National Institute of General Medical Sciences of the National Institutes of Health under award number 1T32GM142623. We acknowledge the Joint Genome Institute (JGI, Proposal ID: 505025) and the Georgia Genomics and Bioinformatics Core (GGBC, UGA, RRID: SCR_010994) for metagenomic sequencing of single-worm isolates and the Georgia Advanced Computing Resource Center (GACRC) at UGA (https://gacrc.uga.edu/) for computational resources that have contributed to the results in this publication.

## Supplementary Figures

**Figure S1. Directory structure and Snakemake rulegraph of MeioBIOME.** The A) organization of the snakemake directory, which includes workflow rules, Conda environments, scripts, and config files, B) structure of the final directory output, and C) Snakemake rulegraph including all the internal subcommands (**Supplementary Table AA**).

## Supplementary Tables

**Table S1. List of all the open-source tools and dependencies implemented in MeioBIOME.** A comprehensive list of the internal subcommands and open-sourced software that are implemented in the Snakemake pipeline. We encourage users to cite the appropriate tools.

**Table S2. Metadata of the publicly available single-worm metagenomic datasets and summary of the host genes recovered.** Nematodes belonging to four nematode families (Thoracostomopsidae, Oncholaimidae, Sphaerolaimidae, and Leptosomatidae) were isolated from four distinct habitats (Tybee Island, Antarctica, Bodega Bay, and Dolphin’s Reef, Florida). The 18S rRNA was recovered using barrnap, and the 18S rRNA gene was BLAST to verify the taxonomic identification of the host. BUSCOs were recovered using the Nematoda-specific OrthoDB v12. The total MAGs include high, medium, and low-quality MAGs.

**Table S3. Summary of the nematode-associated bacterial MAGs recovered using MeioBIOME.** The colors indicate whether the metagenome-assembled genomes (MAGs) were classified as high (green), medium (blue), or low (light red) quality according the CheckM2. The two contaminant MAGs are highlighted dark red. MAGs were classified using the GTDB-Tk workflow.

