## Supplementary figures and images for "MeioBIOME: A snakemake workflow for the parallel analysis of meiofaunal genomes and host-associated bacteria/archaea"

### Figure S1

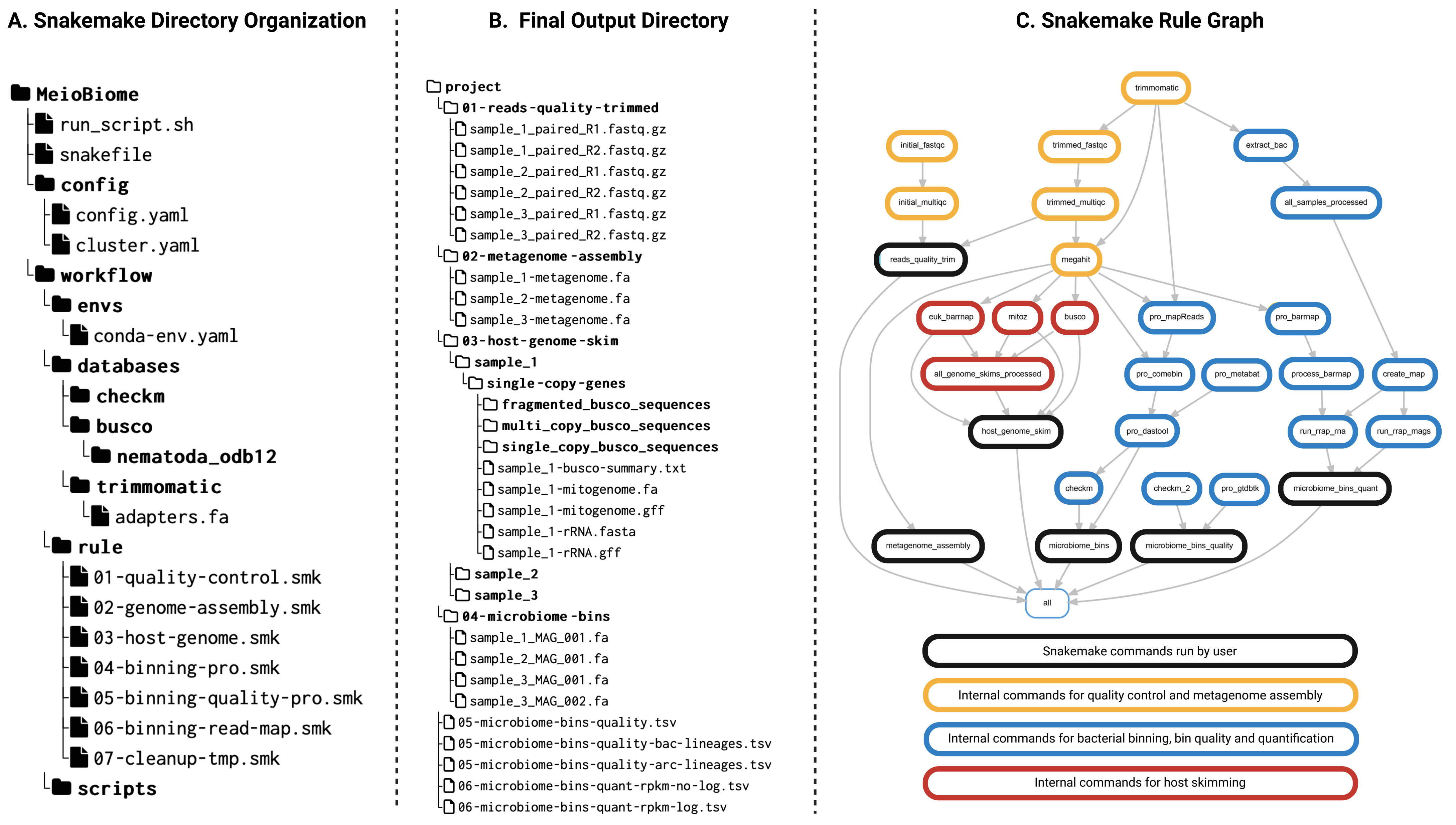
